# Accurate and Rapid Prediction of Protein p*K*_a_: Protein Language Models Reveal the Sequence-p*K*_a_ Relationship

**DOI:** 10.1101/2024.09.16.613101

**Authors:** Shijie Xu, Akira Onoda

## Abstract

Protein p*K*_a_ prediction is a key challenge in computational biology. In this study, we present pKALM, a novel deep learning-based method for high-throughput protein p*K*_a_ prediction. pKALM uses a protein language model (PLM) to capture the complex sequence-structure relationship of proteins. While traditionally considered a structure-based problem, our results show that a PLM pre-trained on large-scale protein sequence databases can effectively learn this relationship and achieve state-of-the-art performance. pKALM accurately predicts the p*K*_a_ values of six residues (Asp, Glu, His, Lys, Cys, and Tyr) and two termini with high precision and efficiency. It excels at predicting both exposed and buried residues, which often deviate from standard p*K*_a_ values measured in solvent. We demonstrate a novel finding that predicted protein isoelectric points (pI) can be used to improve the accuracy of p*K*_a_ prediction. High-throughput p*K*_a_ prediction of the human proteome using pKALM achieves a speed of 4,965 p*K*_a_ predictions per second, which is several orders of magnitude faster than existing state-of-the-art methods. The case studies illustrate the efficacy of pKALM in estimating p*K*_a_ values and the constraints of the method. pKALM will thus be a valuable tool for researchers in the fields of biochemistry, biophysics, and drug design.

## Introduction

Protein p*K*_a_ is crucial for understanding various biological processes, including enzyme catalysis,^1–3^ protein folding,^4–6^ and ligand binding.^7,8^ Chemical groups in the side chain and terminals can be protonated or deprotonated depending on the pH of the environment, affecting protein structure, stability, and function. Currently, the most reliable p*K*_a_ values of proteins are only obtained through laborious, costly, and challenging NMR titration experiments.^9^ Computational methods for the prediction of protein p*K*_a_ have been developed to provide relatively fast and convenient p*K*_a_ estimations while maintaining reasonable accuracy. One widely used method is the empirical approach, PROPKA,^10,11^ which employs a set of human-designed rules to predict p*K*_a_ values based on a single protein structure. PROPKA is considered one of the fastest in protein p*K*_a_ prediction due to the efficient computation, though its accuracy is limited by the simplicity of its rules. In contrast, constant pH molecular dynamics (CpHMD) simulation methods,^12–16^ based on the molecular dynamics (MD) simulations, are considered to be the most accurate computational methods for p*K*_a_ prediction. However, they require a long-time sampling of different protein conformations and protonation states, making them computationally intensive and unsuitable for high-throughput p*K*_a_ prediction. Furthermore, Poisson-Boltzmann (PB) methods,^17–19^ based on continuum electrostatics theory,^20^ provide the p*K*_a_ values for ionizable residues in proteins. PB methods reduce computational costs compared to CpHMD simulations, though at the expense of accuracy. This accuracy decreases due to theoretical assumptions on the basis of modeling the protein-solvent interface as a continuum medium^13^ and single protein conformations in the calculations.

Protein p*K*_a_ prediction is typically approached as a structure-based problem, where the protein structure serves as input features of predcitors. However, several issues arise with this approach, especially for high-throughout prediction: 1) Protein structures are not always available for all proteins, limiting the applicability of structure-based methods. Although structure predictors such as AlphaFold2^21^ and RoseTTAFold^22^ can provide highly accurate atomic-level structures, their high computational cost makes them unsuitable for high-throughput p*K*_a_ prediction. Furthermore, studies indicate that the p*K*_a_ prediction based on structures predicted by AlphaFold2 tend to underestimate of the p*K*_a_ shifts,^23^ rending predictions for significantly shifted p*K*_a_ values less reliable. 2) Protein structures are inherently flexible and dynamic, and specific regions can be ambiguous,^24^ affecting prediction accuracy. Some p*K*_a_ predictors use clustering of different conformations to enhance prediction accuracy.^25^ However, this clustering process is time-consuming and does not fully address the aforementioned structural discrepancy issue. These discrepancies between structures^26^ pose significant challenges to the prediction accuracy of the existing methods. These challenges highlight the need for improved methods in protein p*K*_a_ prediction that can account for structural discrepancies, flexibility, and the availability of protein structures.

Machine learning (ML) methods have recently emerged as a promising approach for protein p*K*_a_ prediction. ML models such as random forests,^27^ graph neural networks,^28^ and multilayer perceptron,^29^ have been applied to predict p*K*_a_ values based on protein structures.^25,30–32^ These ML methods are built with numerous parameters and trained on data sets consisting experimentally determined p*K*_a_ values^33,34^ or p*K*_a_ values derived from other computational methods.^23,35^ Due to their flexible architectures and scalability, these methods effectively model the complex relationship between the protein structures and p*K*_a_ values of protein residues. ML methods achieve both high efficiency and high accuracy in p*K*_a_ prediction compared to traditional methods.

In recent years, large language models (LLMs) have achieved significant success in various modeling tasks, including natural language processing (NLP) and computer vision (CV). Analogously, protein language models (PLMs) have been developed using architectures such as transformers^36,37^ and the recurrent neural network (RNN).^38^ In a PLM, each residue in a sequence is represented as a high-dimensional vector encapsulating the essential information of the sequences and its context. For instance, residues in similar protein environments are likely represented by proximate vectors in the high-dimensional space. PLMs have demonstrated their effectiveness in various protein property prediction tasks, including secondary structure prediction,^39^ contact prediction,,^36^ atomic-level structure prediction,^37^ and intrinsic disorder prediction,^40^ in terms of prediction accuracy and computational efficiency.

This study introduces pKALM, a novel protein p*K*_a_ prediction method based on a protein language model. pKALM is a deep learning-based approach capable of predicting p*K*_a_ values solely from protein sequences. It has been trained to predict the p*K*_a_ values of the side chains of Asp, Glu, His, Lys, Cys, Tyr, as well as the N-terminus and C-terminus of proteins. The performance of pKALM was evaluated across a range of residue types, p*K*_a_ shifts and positions within proteins. The results demonstrated that pKALM outperformed existing methods. Two isoelectric point (pI) predictors were also developed, achieving state-of-the-art performance, and integrated into the pKALM framework to enhance p*K*_a_ prediction accuracy. Furthermore, we demonstrate the high-throughput prediction of p*K*_a_ values for all proteins in the human proteome, emphasizing the significantly faster prediction capabilities of pKALM. A case study is also provided to elucidate the reasons why sequence-based pKALM can effectively predict p*K*_a_ values, with the interpretability of the visualized attention maps of the PLM.

## Method

### Data sets

We utilized the recently released PKAD-2 data set^34^ to train and test various protein p*K*_a_ prediction methods. PKAD-2 compiles experimental p*K*_a_ data from multiple references, providing 1,456 p*K*_a_ values for wild-type proteins and 269 p*K*_a_ values for mutant proteins.

We meticulously revised the original data set to correct missing or erroneous information, excluding unverifiable or non-numerical p*K*_a_ values. The revised data set now includes 1,450 p*K*_a_ values for 165 wild-type proteins and 262 p*K*_a_ values for 47 mutant proteins. The revised PKAD-2 is available in the associated GitHub repository https://github.com/xu-shi-jie/pKALM.

The PKAD-2 data set exhibits considerable redundancy, with homologous proteins having very similar p*K*_a_ values for the same residues. This redundancy presents two critical issues: 1) it causes overlap between training and test sets, compromising the fairness of ML model evaluations, and 2) it hampers the generalization of ML models to novel, non-homologous proteins. To address these issues, we utilized the cd-hit program^41^ to cluster the revised PKAD-2 data set at 30% sequence similarity, a critical threshold (compared to 30% in the work of Cai *et al.*^31^ and 90% in the work of Reis *et al.*^32^). The resulting non-redundant data set was randomly divided into training and test sets in a 5:2 ratio, with 376 p*K*_a_ values in the training set and 152 p*K*_a_ values in test set. Notably, the size of test set closely aligns with those used in related studies.^31^

The literature bias results in an extremely unbalanced distribution of p*K*_a_ values among different residues. Predictions for less abundant residues (Cys, Tyr, N-terminus, and C-terminus) are more challenging compared to abundant residues (Asp, Glu, His, and Lys). Table 1 presents the p*K*_a_ values for both abundant and less abundant residues. The detailed data set construction process is described in Supporting Information, and the p*K*_a_ value distribution (Figure S. 1).

**Table 1:**
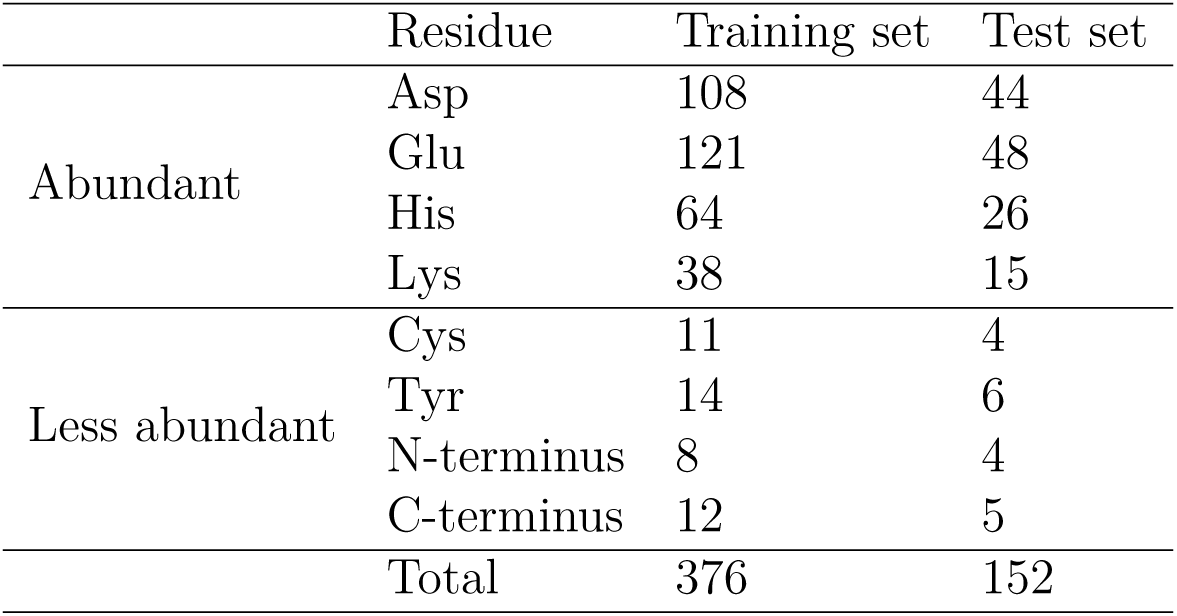
The numbers of p*K*_a_ values of different residues in the data set.

We utilized isoelectric point (pI) data from Kozlowski’s work,^42^ which includes both peptide and protein pI data sets. These data sets, derived from references containing experimental pI values, comprise 119,092 peptide pI values and 2,324 protein pI values. Redundancies in both data sets were removed, and they were split into training and test sets in a 3:1 ratio. Figure S. 2 illustrates the detailed distribution of pI values. Notably, compared to the peptide pI data, the protein pI data are significantly fewer and more concentrated, posing a

### Protein language model

Protein language models are large-scale neural networks pre-trained on extensive unlabeled protein sequences. Architectures such as to BERT^43^ or T5^44^ are commonly employed for PLMs, characterized by parameters counts ranging from millions to billions, each exhibiting distinct properties. In this study, we evaluated nine PLMs of varied architectures and sizes, including the ESM series^36,37^ and ProtTrans series.^39^ Detailed information are provided in Table S. 1.

The scaling law^45^ of large language models demonstrates that their performance consistently improves with larger model and training data sizes. This finding aligns with previous research,^40^ indicating that performance increases with PLM model size but eventually plateaus due to the limited fine-tuning data. Consequently, selecting appropriate PLMs is critical for tasks involving protein property prediction, contingent upon the scale of available fine-tuning data.

### Architecture of pKALM

The architecture of pKALM predictor is depicted in Figure 1(a). The input comprises the protein sequence, encoded through a protein language model, a protein pI model, and a peptide pI model to derive feature vectors. These models are held frozen during pKALM training. The concatenated feature vectors are then fed into a bi-directional long short-term memory (BiLSTM)^46^ network to capture sequential patterns in the protein sequences. To distinguish the residue types (i.e., Asp, Glu, His, Lys, Cys, Tyr, N-terminus, and C-terminus), they are encoded via an embedding layer and added to the BiLSTM network outputs. The combined vectors are subsequently processed by a simple fully connected layer to predict the p*K*_a_ shift. The final predicted p*K*_a_ is derived from the sum of the p*K*_a_ shift and the standard p*K*_a_, consistent with prior work,^11^ as detailed in Table S. 2.

**Figure 1:**
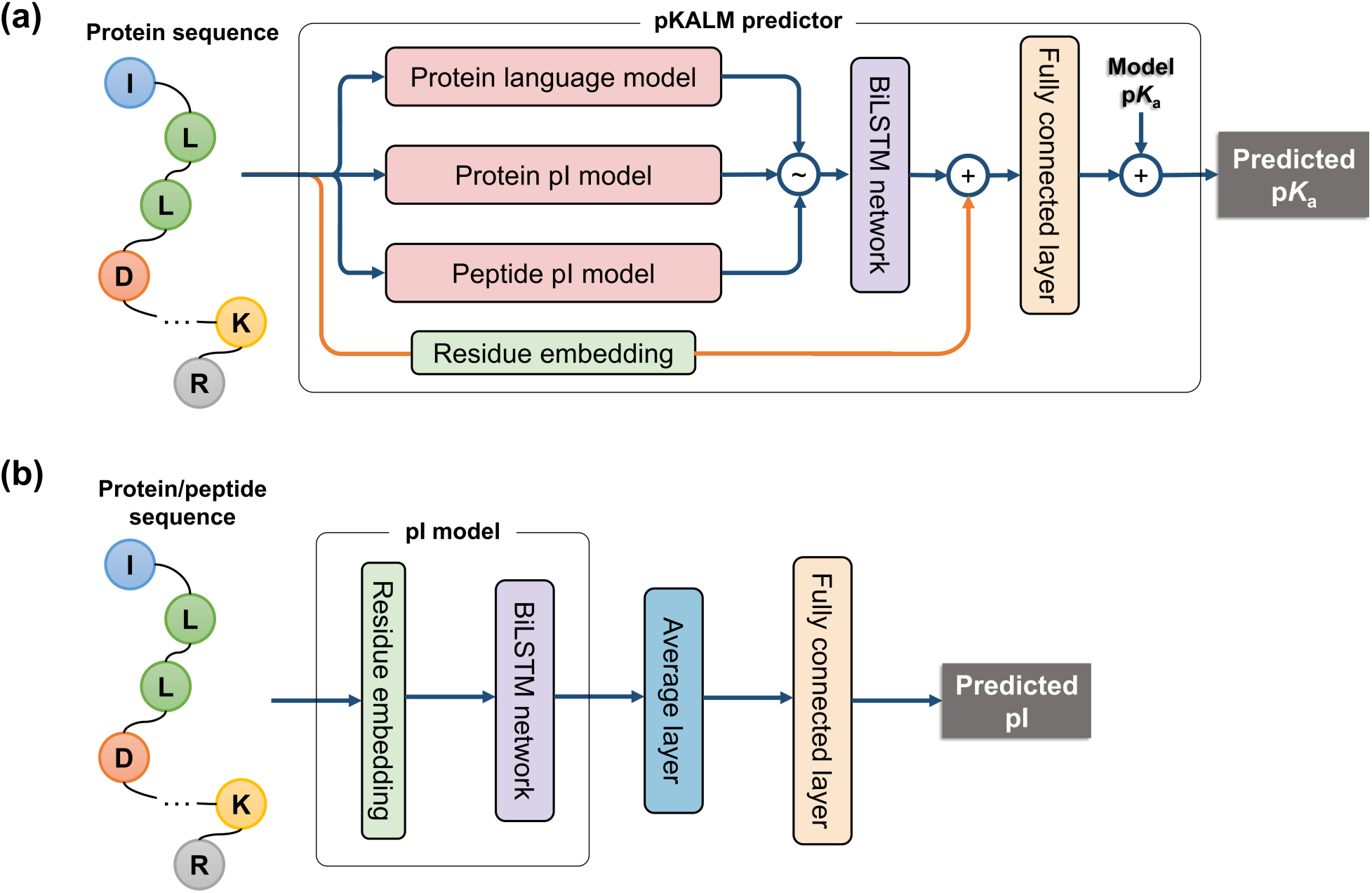
Overview of pKALM. (a) The architecture of pKALM. Arrows indicate data flow, while colored blocks represent different neural networks. The symbol “*∼*” denotes concatenation of feature vectors and “+” denotes vector addition. Flow bifurcation indicates data duplication. (b) Architecture of the pI predictors. The pI model is shared by the peptide pI model and protein pI model.

Furthermore, both pI models share identical architectures, as illustrated in Figure 1(b). The input of each pI model includes either a protein or peptide sequence, encoded into vector sequences via a residue embedding layer. These vectors are then fed into a BiLSTM network to learn sequential patterns. From the BiLSTM outputs, only the hidden states corresponding to protonatable residues (i.e., Arg, Asp, Glu, His, Lys, Cys, and Tyr) and the termini are extracted and averaged to form the final feature vector. A fully connected layer converts this vector into the pI value. Each pI model is trained on distinct data sets: the peptide pI data set and the protein pI data set, respectively.

### Compared methods and data preparation

We compare pKALM with five existing p*K*_a_ prediction methods including PROPKA,^11^ PKAI,^32^ PypKa,^19^ DeepKa,^31^ and PKAI+.^32^ For DeepKa predictions, we utilized the on-line service available at http://www.computbiophys.com/DeepKa/main, while for the other methods, we installed their respective software packages locally and assessed their performance.

As both p*K*_a_ and pI prediction involve regression tasks, we employed root-mean-square error (RMSE) and mean absolute error (MAE) to evaluate method performance. RMSE, preferred for its sensitivity to data set outliers, served as the primary metric. Furthermore, for pI prediction, we utilized RMSE, MAE, and the coefficient of determination (*R*^2^) to comprehensively assess method effectiveness.

All protein structures within our test set were prepared as below. We retrieved PDB files from the Protein Data Bank.^47^ For proteins lacking PDB entries (UniProt IDs: P03230 and P53378), we employed ColabFold^48^ to predict their structures. Repairing the protein structures involved using openMM^49^ to handle non-canonical residues (e.g. selenomethionine is replaced to methionine), mutation of wild-type residues, and removal of HETATM records and water molecules. Missing hydrogen atoms were added using the Amber 14 force field^50^ and Modeller,^51^ assuming a pH of 7.0 and standard p*K*_a_ values for residues.

## Results and discussion

### Hyperparameter tuning and optimization

We utilize a grid search methodology alongside the wandb package^52^ to optimize the hyperparameters of models in the experiments. Specifically, we adjust the dimensions of the hidden states and the layers within the BiLSTM networks for both the pKALM and two pI models. We explore hidden state dimensions and the number of layers in different ranges (see Table S. 3). The optimal hyperparameters are selected based on the performance evaluated through 5-fold cross-validation on the training data set.

We employ the AdamW optimizer^53^ throughout our experiments, with a fixed learning rate of 0.001 and a weight decay of 0.001 to mitigate overfitting. Remaining optimizer hyperparameters are set to their default values. Furthermore, we utilize a cosine annealing learning rate scheduler. The batch sizes are set to 32, with model training spanning 100 epochs for protein p*K*_a_ prediction, 20 epochs for peptide pI prediction, and 10 epochs for protein pI prediction. Loss functions employed are mean-absolute-error (MAE) for p*K*_a_ prediction and mean-squared-error (MSE) for pI prediction.

All experiments are conducted on a Linux workstation equipped with an AMD Ryzen 9 7950X3D 16-core processor, 64 GB memory, and dual NVIDIA RTX 3090Ti GPUs. Implementation utilizes Python 3.12.2 and PyTorch 2.2.1 with CUDA 11.8. For the PLM inference, we utilize Hugging Face’s `transformers` package (ver 4.39.3)^54^ and `accelerate` package (ver 0.28.0).^55^ Except the native automatic mixed-precision feature of PyTorch,^56^ no additional optimization techniques are employed to accelerate the model training and inference.

During the training step of a mini-batch, the protein sequences varied in length were pad and truncated into a fixed length for efficient batch processing. We segment longer sequences into smaller segments with a maximum length of 960 tokens (i.e., residues), due to the constraint that some PLMs, such as ESM1B_650M, only accept sequences shorter than 1024 tokens. To retain contextual information, an additional 31 tokens at the end of the segments are appended to each segment, facilitating overlap between adjacent segments, which will be discarded. Final sequence encodings are derived from the concatenated non-overlapping of segment encodings.

### Prediction of peptide and protein isoelectric point

The isoelectric point (pI) of a protein is the pH at which the protein exhibits no net charge, while the p*K*_a_ values denote the pH at which protein residues are half-protonated. The relationship between the p*K*_a_ values of proteins and its isoelectric point (pI) can be elucidated through the Henderson-Hasselbalch equation for weak acids:^57^

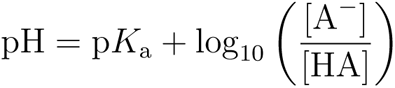

The average charge of an ionizable residue at a specific pH can be calculated by [A*^−^*]*/*[HA] = 10^pH*−*p^*^K^*^a^ . The total charge of the protein can be determined by summing the charges of all ionizable groups. The isoelectric point (pI) is thus the pH at which the total charge of the protein is zero. For instance, the pI of a small molecule with two ionizable groups, with p*K*_a_^1^ and p*K*_a_^2^, can be calculated as pI = (p*K*_a_^1^ + p*K*_a_^2^)*/*2.

Henderson-Hasselbalch equation suggests that pI is closely related to the p*K*_a_ values and this study initially trains peptide and protein pI predictors using certain data sets and integrates them into the pKALM framework. Performance comparisons against state-of-the-art methods for peptide pI prediction are summarized in Table 2. Our peptide pI predictor exhibits superior performance in terms of RMSE, MAE, and *R*^2^ compared to IPC2, the previous leading method trained on identical data sets. 5-fold cross-validation results for peptide pI prediction are illustrated in Figure S. 4, with the optimal hyperparameters selected.

**Table 2:** Performance comparison of different methods for peptide isoelectric point prediction. Bold font indicates our method.

| Method | RMSE | MAE | $R^2$ |
| --- | --- | --- | --- |
| **pKALM, this work** | **0.2204** | **0.1071** | **0.9765** |
| IPC2<sup>42</sup> | 0.2216 | 0.1216 | 0.9761 |
| Bjellqvist<sup>58</sup> | 0.4051 | 0.2836 | 0.9204 |
| Nozaki<sup>59</sup> | 0.4083 | 0.2673 | 0.9191 |
| DTASelect<sup>60</sup> | 0.4235 | 0.2796 | 0.9130 |
| Thurlkill<sup>61</sup> | 0.4466 | 0.2535 | 0.9033 |
| Sillero<sup>62</sup> | 0.4747 | 0.2696 | 0.8907 |
| Dawson<sup>63</sup> | 0.4910 | 0.2642 | 0.8831 |
| Wikipedia<sup>a</sup> | 0.5178 | 0.2974 | 0.8700 |
| Grimsley<sup>64</sup> | 0.5264 | 0.3796 | 0.8656 |

<sup>a</sup>The Wikipedia method refers to the $pK_a$ values as presented in the Wikipedia page in 2005.

We develop a protein pI predictor and evaluate its performance against existing benchmarks. Performance comparisons against state-of-the-art methods for protein pI prediction are summarized in Table 3. Our method achieves the third-best RMSE and *R*^2^ scores, and the fourth-best MAE among the methods evaluated. Notably, our approach employs deep learning, which typically demands larger data sets for excellent performance. The limited size of available protein pI data sets (see Section “Data set”) constrains the efficacy of our predictor despite employ a comparable architecture. 5-fold cross-validation results for protein pI prediction are depicted in Figure S. 5, with the optimal hyperparameters selected.

**Table 3:** Performance comparison of different methods for protein isoelectric point prediction. Bold font indicates our method.

| Method | RMSE | MAE | $R^2$ |
| --- | --- | --- | --- |
| IPC2<sup>42</sup> | 0.8479 | 0.5906 | 0.5934 |
| IPC<sup>65</sup> | 0.8677 | 0.6109 | 0.5760 |
| **pKALM, this work** | **0.9069** | **0.6468** | **0.5589** |
| ProMoST<sup>66</sup> | 0.9113 | 0.6444 | 0.5183 |
| Toseland<sup>67</sup> | 0.9278 | 0.6537 | 0.5095 |
| Dawson<sup>63</sup> | 0.9365 | 0.6586 | 0.4977 |
| Bjellqvist<sup>58</sup> | 0.9369 | 0.6536 | 0.5005 |
| Wikipedia<sup>a</sup> | 0.9484 | 0.6795 | 0.4860 |
| Rodwell<sup>68</sup> | 0.9579 | 0.6762 | 0.4706 |
| Grimsley<sup>64</sup> | 0.9588 | 0.6953 | 0.4779 |

<sup>a</sup>The Wikipedia method refers to the $pK_a$ values as presented in the Wikipedia page in 2005.

### Performance of different protein language models

We assess the performance of multiple protein language models for predicting p*K*_a_ values, as depicted in the blue and green curves in Figure 2. The evaluation includes validation and test RMSE for nine PLMs: eight PLM-based configurations (with “ESM” or “PT” prefixes) and a compared configuration replacing PLM with an embedding layer (“No PLM”). The points in the curves representing the best configurations selected from validated RMSEs, which consistently decreases with an increase in model parameters from 8 to 650 millions, and eventually achieve the lowest for three largest PLMs: ESM2_650M, ESM2_3B, and ESM2_15B. Other PLMs (ESM1B_650M, PT_BFD, PT_UR), and the non-PLM configuration exhibit varying performances due to their distinct architectures and training data sets from ESM2 series, and their details can be found in the original literature.^36,39^

**Figure 2:**
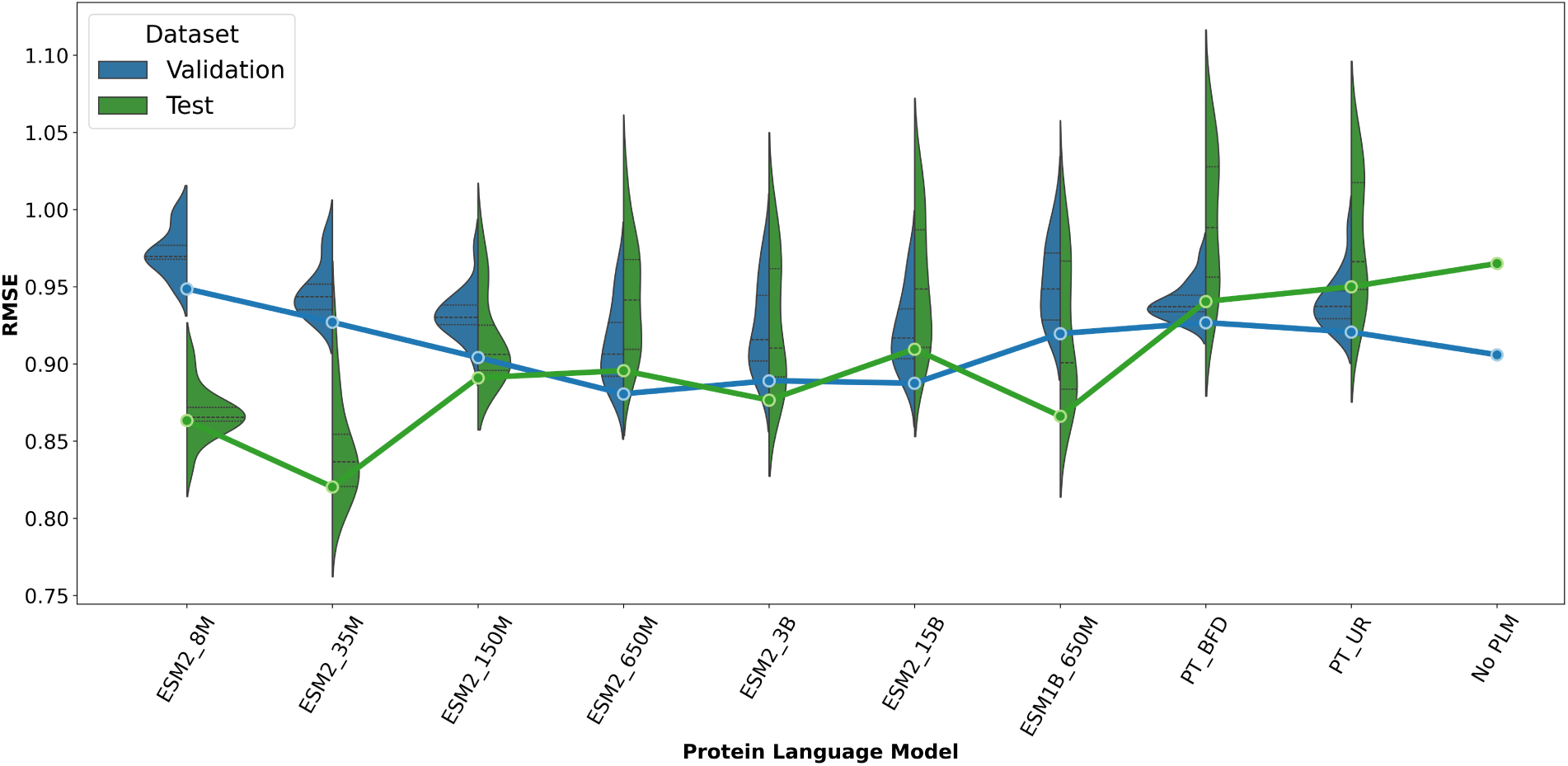
Comparison of the performance of different protein language models on the validation and test sets for all models and the best model selection. The violin graph illustrates the distribution of (Validation/Test) RMSEs for different PLMs with different hyperparameters (hidden states and layers). The curves represent the best (Validation/Test) RMSEs selected based on the validation set. Note that for the non-PLM configuration, the distribution of RMSEs across different configurations is very concentrated and we only show the best RMSE.

The test RMSEs, however, diverges notably from the validation RMSEs. We define the p*K*_a_ shift as the difference between model-predicted and experimentally determined p*K*_a_, categorizing shifts into five ranges split by 0.5 pH units: 0-0.5, 0.5-1.0, 1.0-1.5, 1.5-2.0, and *>*2.0. Figure S. 3 illustrates the test RMSE of different PLMs across these ranges. Notably, larger PLMs (ESM1B_650M, ESM2_3B, and ESM2_15B) exhibits superior performance when p*K*_a_ shifts are small (<1.0 pH unit). In contrast, for shifts larger than 1.5, the configurations with smaller PLMs (ESM2_8M and ESM2_35M), or without PLM, demonstrate better performance. These findings suggest that larger PLMs excel in capturing evolutionary information from protein sequences and tend to predict normal p*K*_a_ values more accurately, while they may be less reliable for p*K*_a_ values that shift significantly. Thus, selecting an appropriate PLM is crucial to balance prediction accuracy of p*K*_a_ values that shift significantly and those that do not.

The distribution of RMSEs for small-size PLMs is more concentrated, while the gaps between the validation and test RMSEs are larger. In contrast, large-size PLMs exhibit less concentrated RMSEs with smaller gaps between validation and test RMSEs, as shown in the violin graph in Figure 2. Overall, the variance in performance across different PLMs and hyperparameters underscores the importance of selecting an appropriate PLM for protein p*K*_a_ prediction and highlights the necessity of hyperparameter tuning for optimal performance. Notably, our choice of PLM is considered *a priori* knowledge rather than a hyperparameter. For subsequent experiments, we prioritize the RMSE of four abundant residues (Asp, Glu, His, and Lys) as the primary metric for selecting the best PLM. ESM2_35M achieves the best performance in terms of test RMSE among the evaluated models.

### pKALM outperforms existing methods

We evaluate the performance of our pKALM method against reported method (Figure 3a and 3b). The results demonstrate that all methods can predict p*K*_a_ values for Asp, Glu, His, and Lys. Importantly, pKALM achieved the lowest test RMSE on four abundant residues (Asp 0.6929, Glu 0.6641, His 1.0468, Lys 0.7701). PKAI+, another deep learning-based method, showed the second-best performance, followed by DeepKa, PypKa, and PKAI. The empirical method PROPKA performs the worst, due to its simplistic empirical rules and limited model complexity. The predictions of Glu residues exhibited relatively small test RMSE among all predictors due to the concentrated p*K*_a_ values of glutamic acid. In contrast, the predictions of His residues had the highest RMSE, due to the limited and diverse distribution of p*K*_a_ values in the data set. Asp residues, with the second largest data (108/44 in training/test set) and a broader p*K*_a_ distribution, provided a fair and challenging to benchmark. Performance comparisons on different residues in terms of RMSE and MAE can be found with details in Table S. 4 and S. 5. Overall, pKALM significant outperformed other methods.

**Figure 3:**
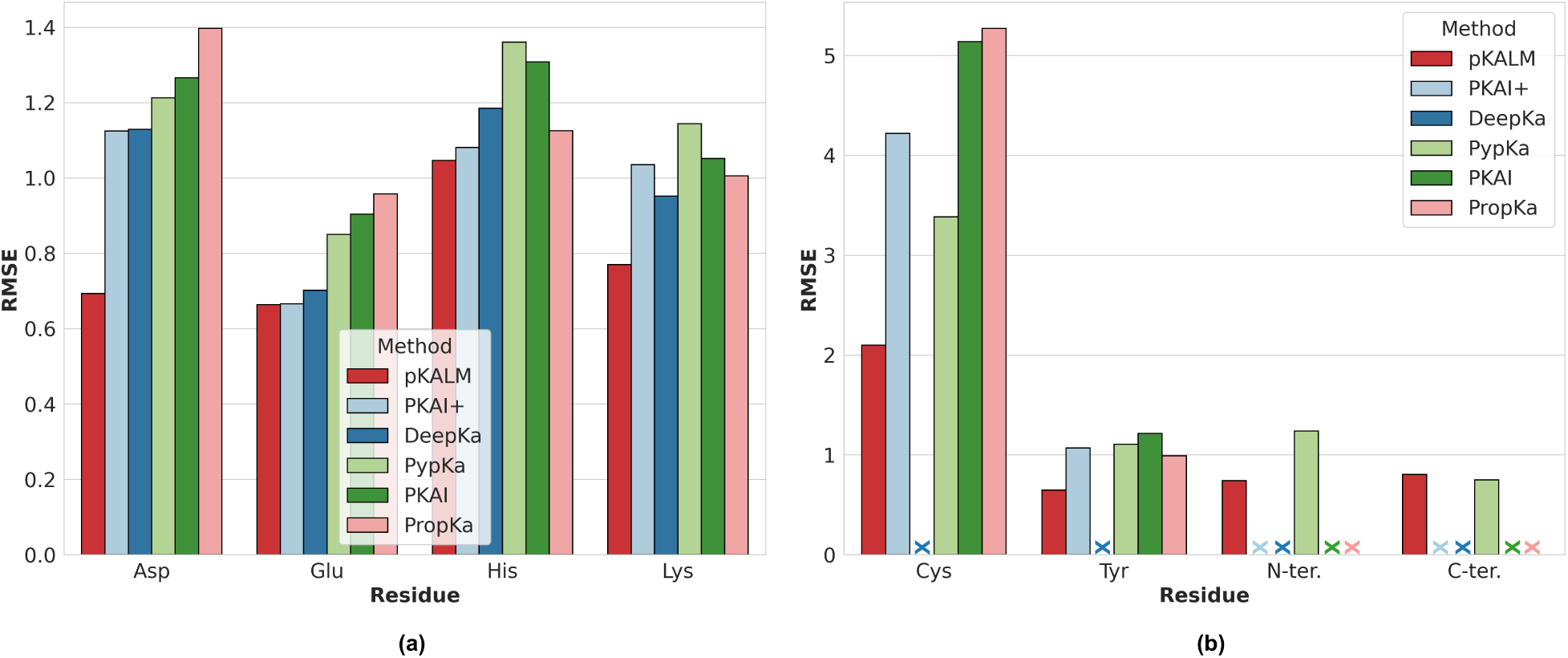
Performance comparison of pKALM with existing methods for (a) abundant residues: Asp, Glu, His, and Lys and (b) less abundant residues: Cys, Tyr, N-terminus (N-ter.), and C-terminus (C-ter.). Some methods are not available for certain residues, as indicated by “X”.

The p*K*_a_ values of less abundant residues Cys, Tyr, N-terminus, and C-terminus were predicted in superior performance in terms of RMSEs (Cys 2.098, Tyr 0.6479, N-terminus 0.7419, C-terminus 0.8038) by pKALM relative to existing methods, as illustrated in Figure 3b. Predictions for Cys were particularly challenging due to the their limited data and very diverse p*K*_a_ values.

We compared pKALM with the available methods, marking unavailable ones with an “X”. pKALM is unique in predicting p*K*_a_ values for Cys with a RMSE of approximately 2 pH units, whereas other methods showed significantly large RMSEs (PKAI+>4, PypKa>3, PKAI and PROPKA>5), rending their predictions for Cys unreliable. Despite a limited test set, pKALM achieved the lowest RMSE for Tyr (14/6 in training/test set) and N-terminus (8/4 in training/test set). Detailed performance comparisons in terms of RMSE and MAE can be found in Table S. 6 and S. 7. pKALM and PypKa provided predictions for C-terminus with comparable accuracy (pKALM 0.8038 vs PypKa 0.7496). This indicates that not-machine-learning-based methods remain dependable for residues lacking sufficient data to train machine learning models.

### Performance on p*K*_a_ shifts

It is crucial to evaluate the performance of p*K*_a_ predictors on different p*K*_a_ shifts. According to the PKAD-2 dataset, most of p*K*_a_ values are within 1 pH unit of their standard values, though some are significantly shifted (e.g., >2 pH units). Predicting small p*K*_a_ shifts is relatively straightforward, while predicting large shifts is more challenging, necessitating evaluation across different p*K*_a_ shift ranges. According to the previous study,^31^ we split the p*K*_a_ shifts into five ranges with 0.5 pH unit interval: 0-0.5, 0.5-1.0, 1.0-1.5, 1.5-2.0 and *>*2.0 and assessed the performance of pKALM and existing methods on these ranges (Figure 4a). pKALM achieves the lowest RMSE in ranges *≤*1.5 while the third best in the range of 1.5-2.0 and the second-best in the range of *>*2.0. It is worth noting that, another two deep learning-based methods, PKAI and PKAI+, also lag behind the best performance for larger ranges. Deep learning methods are trained for minimizing the overall loss function and may not always focus on learning the largely shifted p*K*_a_ values. Considering the large part of small p*K*_a_ shifts in the data set, it is reasonable that the deep learning methods are not the best for the largely shifted p*K*_a_ values. On the other hand, DeepKa is optimized for relatively balanced distribution of p*K*_a_ shifts and thus performs better than PKAI and PKAI+ for the largely shifted p*K*_a_ values while still worse than pKALM in terms of accuracy. PROPKA is the best one for the p*K*_a_ shifts larger than 2.0 pH units while the data point of this range is very limited. Furthermore, its performance on small p*K*_a_ shifts is relatively low, suggesting such a method not reliable for the general p*K*_a_ prediction since we do not know which range the p*K*_a_ value belongs to before the prediction. In contrast, our method pKALM achieves balanced performance across different p*K*_a_ shift ranges.

**Figure 4:**
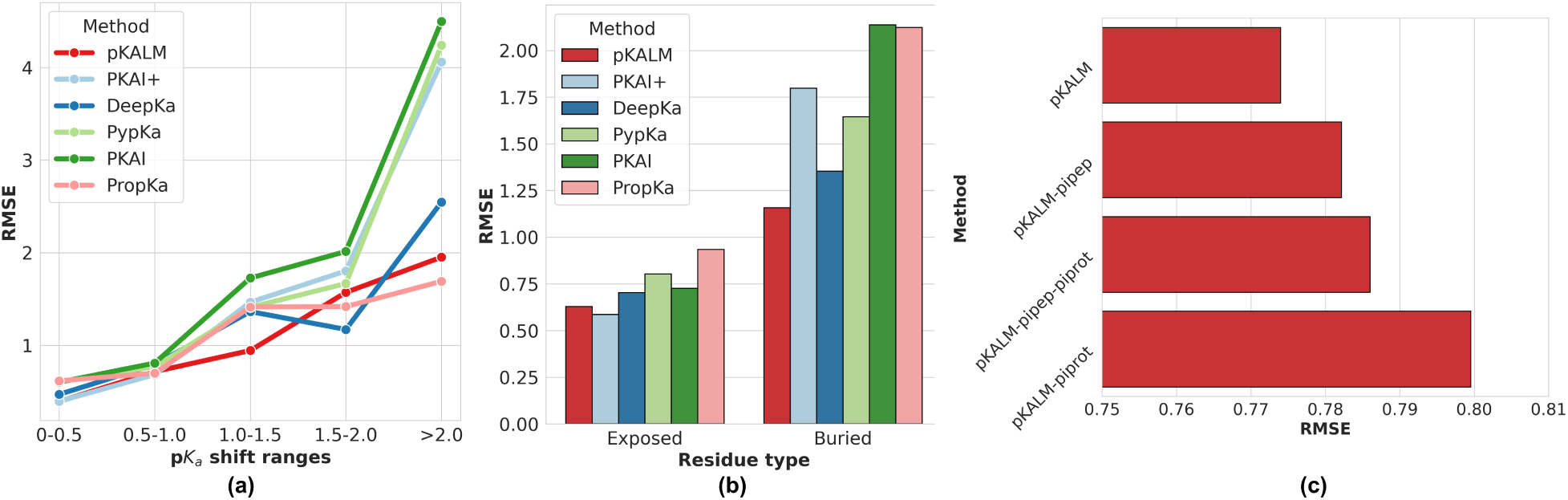
(a) Performance comparison of pKALM with reported methods for largely shifted p*K*_a_ values. (b) Performance comparison of pKALM with reported methods for exposed and buried residues. (c) Ablation study of pKALM by removing the pI models.

### Performance on the exposed and buried residues

We evaluate the performance of reported methods for exposed and buried residues separately. The definition of exposed and buried residues are based on the relative solvent accessibility (RSA) of the residues, which is already calculated in the PKAD-2 data set. We choose 50% as the threshold, same to the previous study^30^ (4b). It is clear that pKALM achieves the lowest RMSE for challenging buried residues. For exposed residues, pKALM also achieves the second-lowest RMSE (0.6281), following PKAI+ (0.5862), while all predictors have relatively similar performance. This is reasonable since the exposed residues are more accessible to the solvent and thus their p*K*_a_ values are more close to the standard p*K*_a_ values, suggesting them much easier to predict. The slight difference between pKALM and PKAI+ may be due to the measurement uncertainty of p*K*_a_ values in the data set, which is usually around (0.01-0.1).

### Ablation study and comparison with existing methods

The necessity of adopting the pI models in p*K*_a_ prediction was confirmed by an ablation study of different modules (Figure 4c). To investigate the importance of modules inside a deep learning model, we remove the pI models one by one, retrain them with the same data sets, and evaluate them on the same test sets. Four versions of configurations are tested: 1) pKALM, 2) pKALM without peptide pI model “pKALM-pipep”, 3) pKALM without protein pI model “pKALM-piprot”, and 4) pKALM without both pI models “pKALM-pipep-piprot”. We observe the changes of the test RMSE of abundant residues and the results show that the pI models are crucial for the p*K*_a_ prediction, though the version without any pI models are good. By removing the peptide pI model, the RMSE increases; removing the protein pI model, the RMSE also increases. When we remove only the protein pI model, the RMSE increases the most. This suggest that protein pI model is the most crucial for this architecture.

### High-throughput prediction

To evaluate the speeds of pKALM, the p*K*_a_ values of all proteins in the human proteome are predicted. The human proteome data set is download from the UniProt website (https://www.uniprot.org/proteomes/UP000005640) which contains a total of 20,598 proteins and 3,554,768 protonatable residues. The whole prediction process took 11 minutes 56 seconds on our machine, which produced an average speed of 4,965 p*K*_a_ per second. The distribution histograms of the predicted p*K*_a_ values of different residues (Figure S. 6) indicates the predictions have a very wide range of p*K*_a_ value distributions for all residues. This suggests that pKALM is capable of predicting the significantly shifted p*K*_a_ values in the proteins. The predicted p*K*_a_ of terminal groups have relatively concentrated distributions, which is consistent with the fact that N-terminal and C-terminal are more likely exposed to the solvent and thus have more concentrated p*K*_a_ to its standard p*K*_a_ values.

We evaluate the speed of different predictors that compared in this study. We cite the reported speeds of DeepKa in the published work.^69^ We have tested the speed on human proteome data set for our pKALM. Other methods are installed with their official programs on our machine. Furthermore, we randomly sample 1,000 structures from the PDB database and predict the p*K*_a_ values of all residues in the structures by PKAI, PKAI+, PROPKA. Since PypKa is very slow, we sampled only 100 structures and predicted their p*K*_a_ values by PypKa. The speed of these predictors are shown in Table 4.

**Table 4:** Speed comparison of different methods for p*K*_a_ prediction.

| Method | Total $pK_a$ | Total time | $pK_a$ per sec |
| --- | --- | --- | --- |
| **pKALM, this work** | **3,554,768** | **11 min 56 s** | **4,964.76** |
| PKAI | 187,312 | 23 min 31 s | 132.75 |
| PKAI+ | 187,312 | 23 min 38 s | 132.10 |
| PROPKA | 219,198 | 96 min 14 s | 37.96 |
| DeepKa | 1,537 | 3 min 35 s | 7.15 |
| PypKa | 11,565 | 3 h 43 min 49 s | 0.86 |

<sup>a</sup> Note that different methods are tested on different data sets.

It is clear that pKALM is the fastest method among the compared methods, which is more than 37 times faster than PKAI/PKAI+, 130 times faster than PROPKA, 694 times faster than DeepKa, and 5773 times faster than PypKa. The speed of pKALM is mainly due to its sequence-based design and the parallel computation of the deep learning models on the GPUs. PLMs are considered as the bottleneck of the deep learning models. However, the p*K*_a_ prediction of pKALM is performed for all protonatable residues in the same protein simultaneously. The price of PLM inference is paid only once for each protein, instead of each residue, which is more efficient than the residue-by-residue prediction. Another deep learning methods, PKAI/PKAI+ also provide the second and the third best speed. Surprisingly, the empirical method PROPKA predicts more slowly than the deep learning methods, possibly due to its implementation on the CPU and the sequential prediction of the residues’ p*K*_a_ values.

### Case study and discussion

#### Prediction for ribonuclease T_1_

We use pKALM to predict the p*K*_a_ values of ribonuclease T_1_ (PDB ID: IRGA) to have a case study. Ribonuclease T_1_ is a small protein consisting of 104 residues (Figure 5a). It has been reported to have p*K*_a_ values as shown in Table 5. The predicted p*K*_a_ values from pKALM are very close to the experimental values. This example demonstrates that pKALM accurately predict the p*K*_a_ values of the residues in the protein.

**Figure 5:**
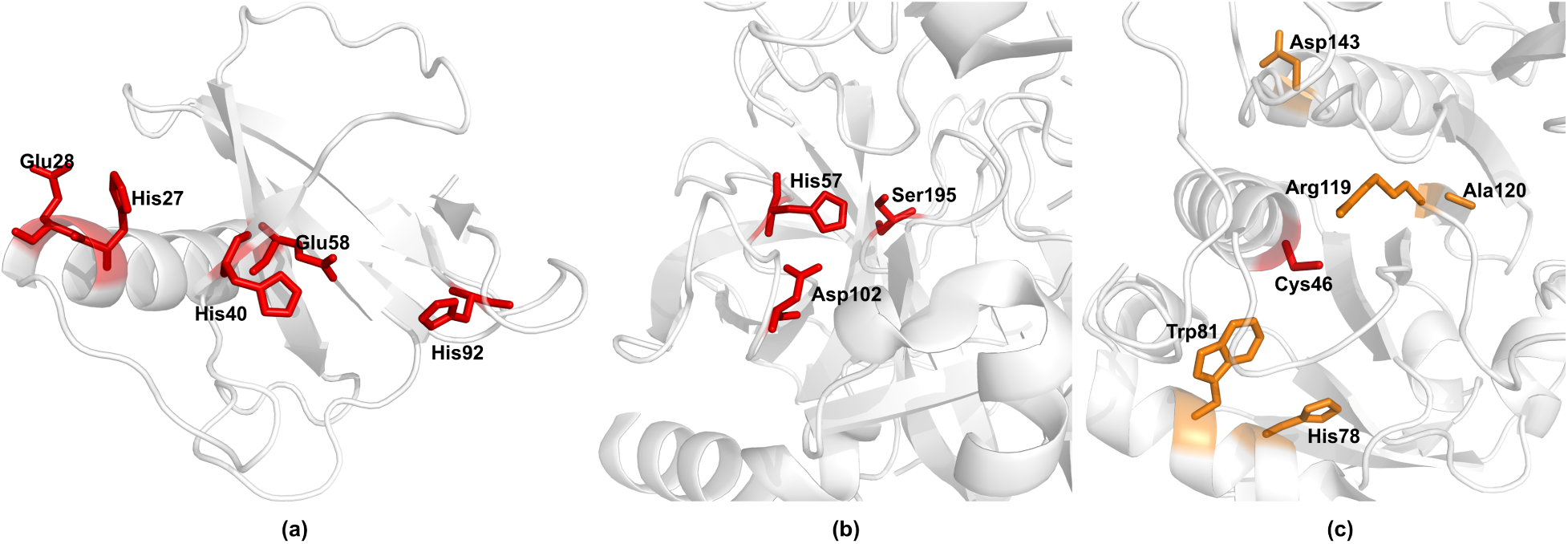
Examplary predictions. (a) The five titrated residues of ribonuclease T_1_. (b) The catalytic triad of chymotrypsin. (c) The bacterial peroxiredoxin AhpC. The catalytic Cys46 is shown in red. Residues with high attention maps to Cys46 are shown in orange.

**Table 5:**
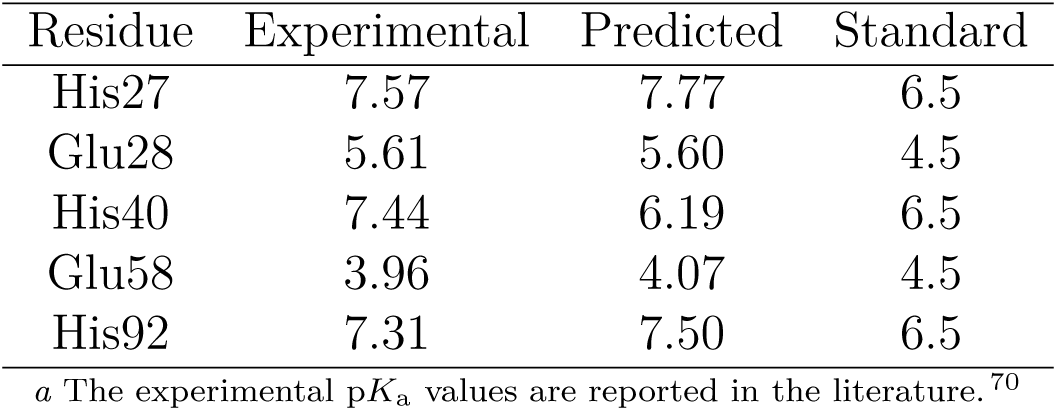
The experimental, predicted, and standard p*K*_a_ values of selected residues in ribonuclease T_1_.

#### Prediction for chymotrypsin

pKALM is employed to predict the catalytic triad motif of chymotrypsin, a serine protease. This motif is comprised of Asp, His, and Ser (Figure 5b). The input for pKALM is the B chain of the apoenzyme structure (PDB ID: 4CHA). Given the strong interaction between these three residues, the p*K*_a_ values of His57 are significantly altered from their standard p*K*_a_ value of 6.5. The predicted p*K*_a_ value of His57 from pKALM is 7.65, which is in close agreement with the experimental value of 7.5.^2^ This indicates that pKALM is effective in capturing the intricate interactions between the residues and accurately predicting the p*K*_a_ values of the residues in the catalytic triad. It is worth noting that Ser in the catalytic triad is not on the input chains but the prediction is still accurate, possibly due to the strong interactions between the residues in the catalytic triad, which are captured by the protein language model.

#### Why PLM-based design works?

We conduct a case study to demonstrate why the PLM-based design of pKALM works very well. We use pKALM to predict the p*K*_a_ values of Cys46 in the bacterial peroxiredoxin AhpC (PDB ID: 4MA9), a large protein consisting of five chains and hundreds of residues. The key residues for analysis are shown in Figure 5c. The p*K*_a_ value of Cys46 is 5.94,^71^ which is largely shifted from the standard p*K*_a_ value of 9.0. The predicted p*K*_a_ value from pKALM is 6.67, which is close to the experimental value, while other methods predict the values including PROPKA (9.0), PKAI (9.64), and PKAI+ (9.08), close to the standard p*K*_a_ value of 9.0. This suggests that pKALM is more reliable for the prediction of the p*K*_a_ values of Cys residues in proteins.

To visualize the effectiveness of pKALM, the attention maps are considered as the useful tool to interpret the deep learning models.^72^ We thus visualize the attention maps of all 20 heads of the PLM ESM2_35M when the input is the sequence of AhpC (Figure S. 7 and Figure S. 8). Most heads only capture the short-distance interactions between the residues, while some heads capture the long-distance interactions. Here we take the 14th head as an example to analyze the attention map, since it has the most significant long-distance interactions. The most important residues that Cys46 pays attention to are His78, Trp81, Arg119, Ala120, and Asp143. These residues are distant from Cys46 in sequence and close in the 3D structure. The attention map imply that pKALM may successfully build the sequence-p*K*_a_ relationship by capturing the long-distance interactions and possibly more essential information of the residues in the protein sequences.

## Conclusions

The understanding of protein functions in biological processes is contingent upon the knowledge of protein p*K*_a_. It is a significant challenge to accurately measure the p*K*_a_ of specific residues in a protein. However, computational methods provide an alternative approach to predict the p*K*_a_ values. In this study, we propose a deep learning-based method, pKALM, which employs a protein language model to achieve highly accurate predictions for Asp, Glu, His, Lys, Cys, Tyr, N-terminus, and C-terminus. Furthermore, two pI models were trained and integrated into the pKALM architecture. The pI models are of particular importance for the prediction of p*K*_a_, with the protein pI model being especially crucial in this regard. The performance of pKALM was evaluated on the PKAD-2 dataset, and it was compared with existing methods. The results demonstrate that pKALM outperforms existing methods in terms of root mean square error (RMSE) and mean absolute error (MAE). Furthermore, the performance of pKALM was evaluated on data sets with significantly shifted p*K*_a_ values, exposed and buried residues, and the human proteome. Since the current version of pKALM is designed for single-chain inputs, the development of the multi-chain adaptable version is currently underway.

## Data availability

All data sets used in this study are publicly available at PKAD-2 official website and the revised data set is available at https://github.com/xu-shi-jie/pKALM. The corresponding reproducible source code is publicly available at https://github.com/xu-shi-jie/pKALM. A web server for pKALM is available at https://onodalab.ees.hokudai.ac.jp/pkalm.

## Author Contributions

All authors conceived and designed the study. S.X. developed the method, performed data analysis, and wrote the paper. A.O. supervised the research project and critically reviewed and revised the manuscript. All authors approved the final manuscript.

## Supporting information

Support Information

## Acknowledgements

This work was supported by Hokkaido University DX Doctoral Fellowship (JST SPRING, Grant Number JPMJSP2119) to S.X. and SATREPS Project “Recovering High-Value Bio-products for Sustainable Fisheries in Chile (ReBiS)” funded by JST/JICA (Grant Number JPMJSA2206) and JSPS KAKENHI Grant Number JP24H02213 in Transformative Research Areas (A) JP24A202 Integrated Science of Synthesis by Chemical Structure Reprogramming (SReP) and JP24K01533 to A.O.

