## Supplementary material for "Accurate and Rapid Prediction of Protein p*K*_a_: Protein Language Models Reveal the Sequence-p*K*_a_ Relationship": Support Information

#### Data processing for constructing the training and test data sets

Firstly, we selected all p$K_a$ values in the PKAD-2 dataset for Asp, Cys, Glu, His, Lys, Tyr, C-terminus, and N-terminus residues, including both wild-type and mutant proteins. Protein structures were obtained from the PDBj website `https://pdbj.org/` using the following command:

```
rsync -rlpt -v -z --delete \
    ftp.pdbj.org::ftp\_data/structures/divided/pdb/ ./wwpdb
```

(up to 7/2/2024). All structures collected were then fixed using the PDBFixer tool under the openMM platform.

Secondly, we verified that the residues recorded in the PKAD-2 dataset matched the PDB structures (residue ID, residue name, and chain ID). We checked for missing residues in their structures, marking unknown residues with an “X” symbol. Sequences were extracted based on tHis analysis, rather than directly from the “pdbres” field in the PDB files.

Next, we removed redundancy with the following command:

```
./cdhit/psi-cd-hit/psi-cd-hit.pl -i $in_path -o $db30 \
    -c 0.3 -ce 1e-6 -G 0 -g 1 -para 8 -blp 4
```

and retained only the representative sequences in each cluster for subsequent analysis. The clustered dataset was then split into training, validation, and test sets with a ratio of 4:1:2 using the following Python code:

```python
test_dfs, val_dfs, train_dfs = [], [], []
for res in df['Res Name'].unique():
    res_df = df[df['Res Name'] == res]
    # res_df is the dataframe of the residue data for each residues
    test_dfs.append(res_df[0::7])
    test_dfs.append(res_df[1::7])

    val_dfs.append(res_df[2::7])

    train_dfs.append(res_df[3::7])
    train_dfs.append(res_df[4::7])
    train_dfs.append(res_df[5::7])
    train_dfs.append(res_df[6::7])
test_df = pd.concat(test_dfs)
val_df = pd.concat(val_dfs)
train_df = pd.concat(train_dfs)
```

### Supplementary figures

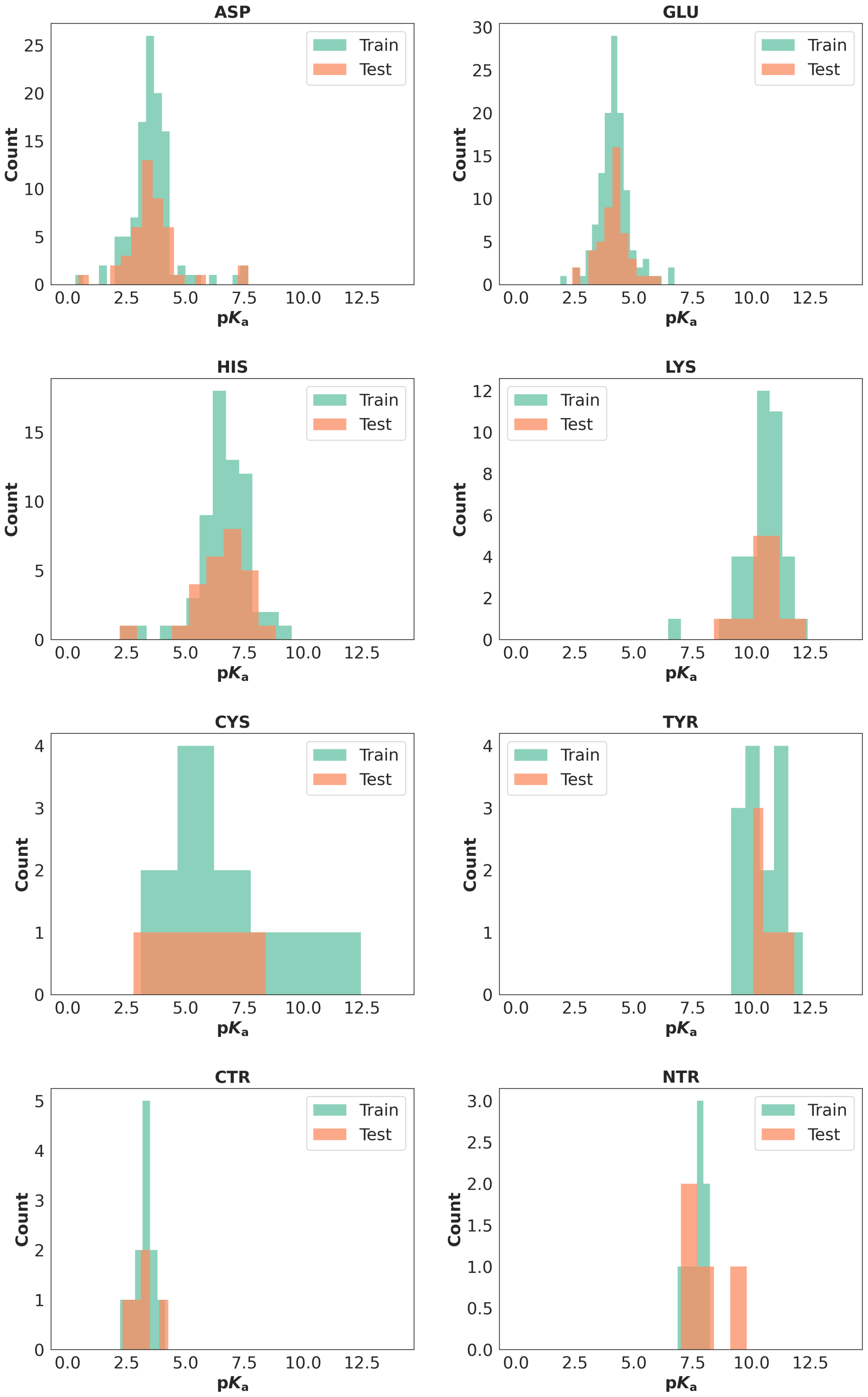


Figure S. 1: The distribution of the p$K_\mathrm{a}$ values in the training and test data sets

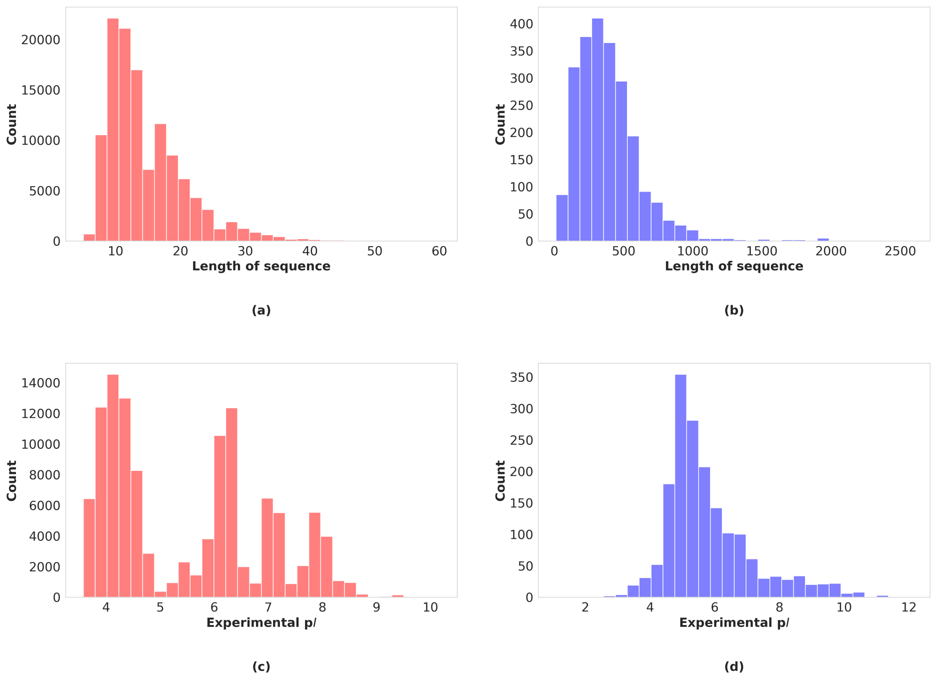


Figure S. 2: (a) The distribution of the sequence lengths of IPC2 peptide data set. (b) The distribution of the sequence lengths of IPC2 protein data set. (c) The distribution of the isoelectric points of IPC2 peptide data set. (d) The distribution of the isoelectric points of IPC2 protein data set.

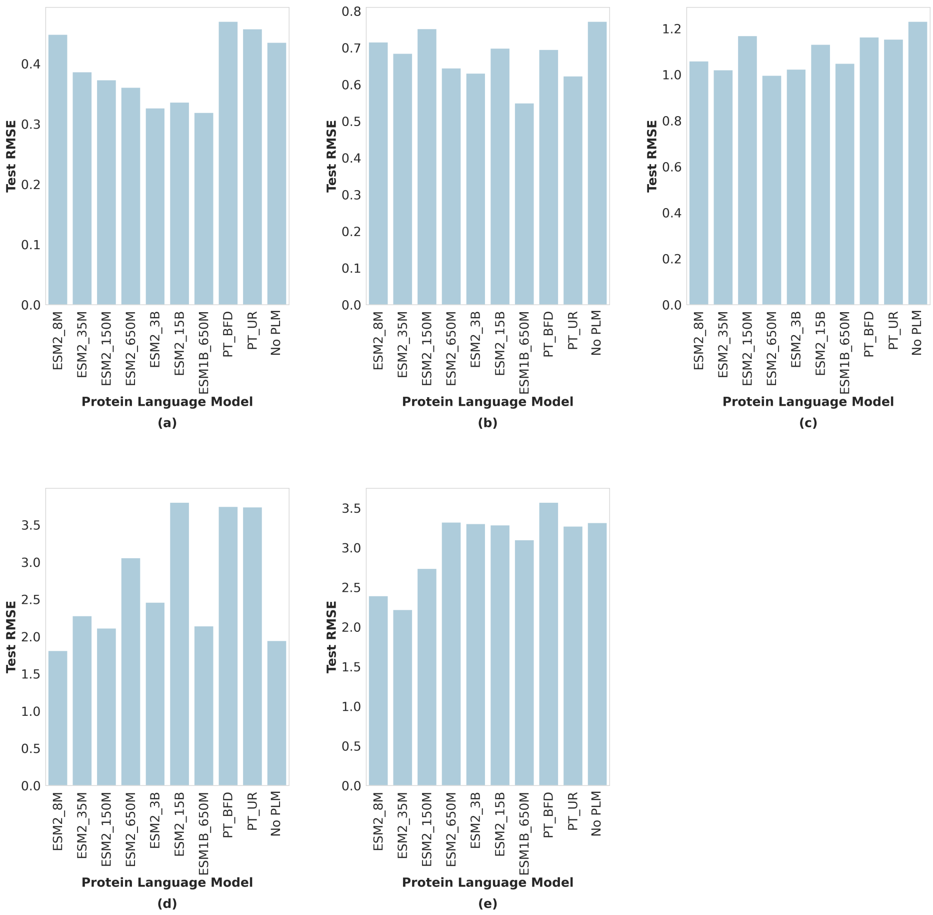


Figure S. 3: Performance of different protein language models on different p$K_{\mathrm{a}}$ shift ranges: (a) 0-0.5, (b) 0.5-1.0, (c) 1.0-1.5, (d) 1.5-2.0, and (e) $> 2.0$

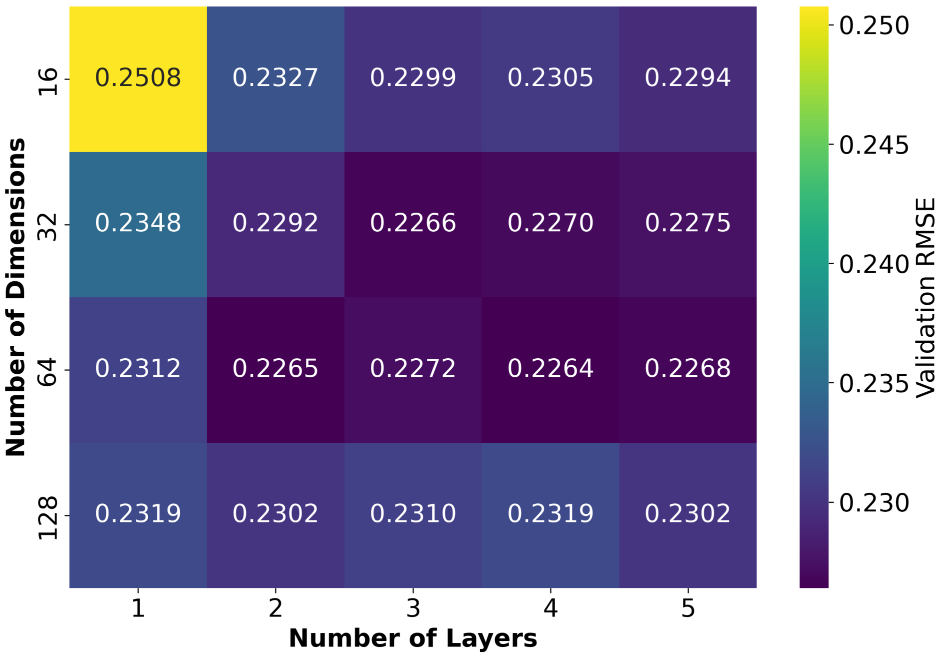


Figure S. 4: Heatmap of validated RMSE of peptide isoelectric point prediction with different hyperparameters

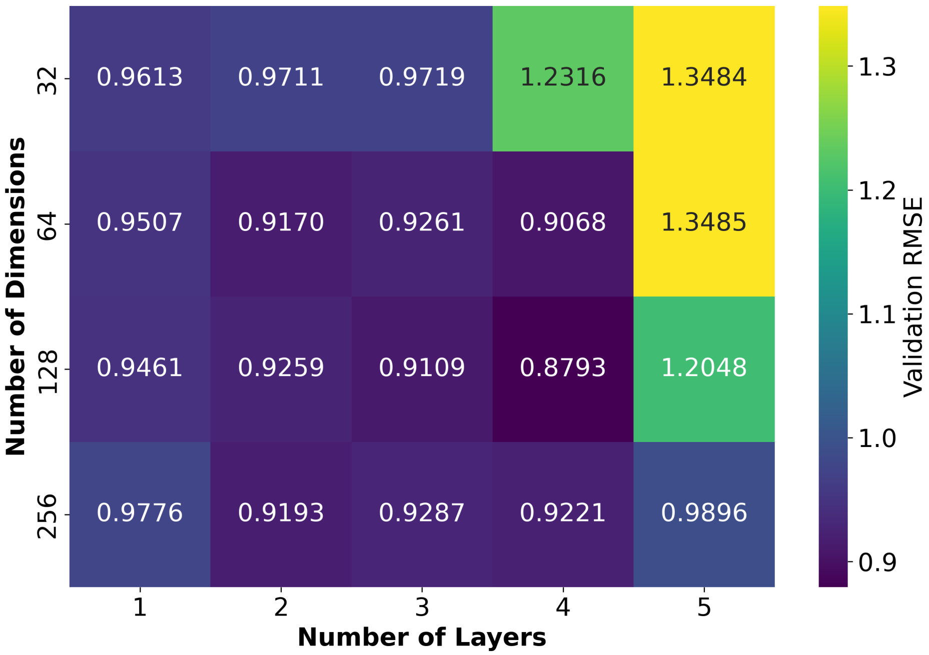


Figure S. 5: Heatmap of validated RMSE of protein isoelectric point prediction with different hyperparameters

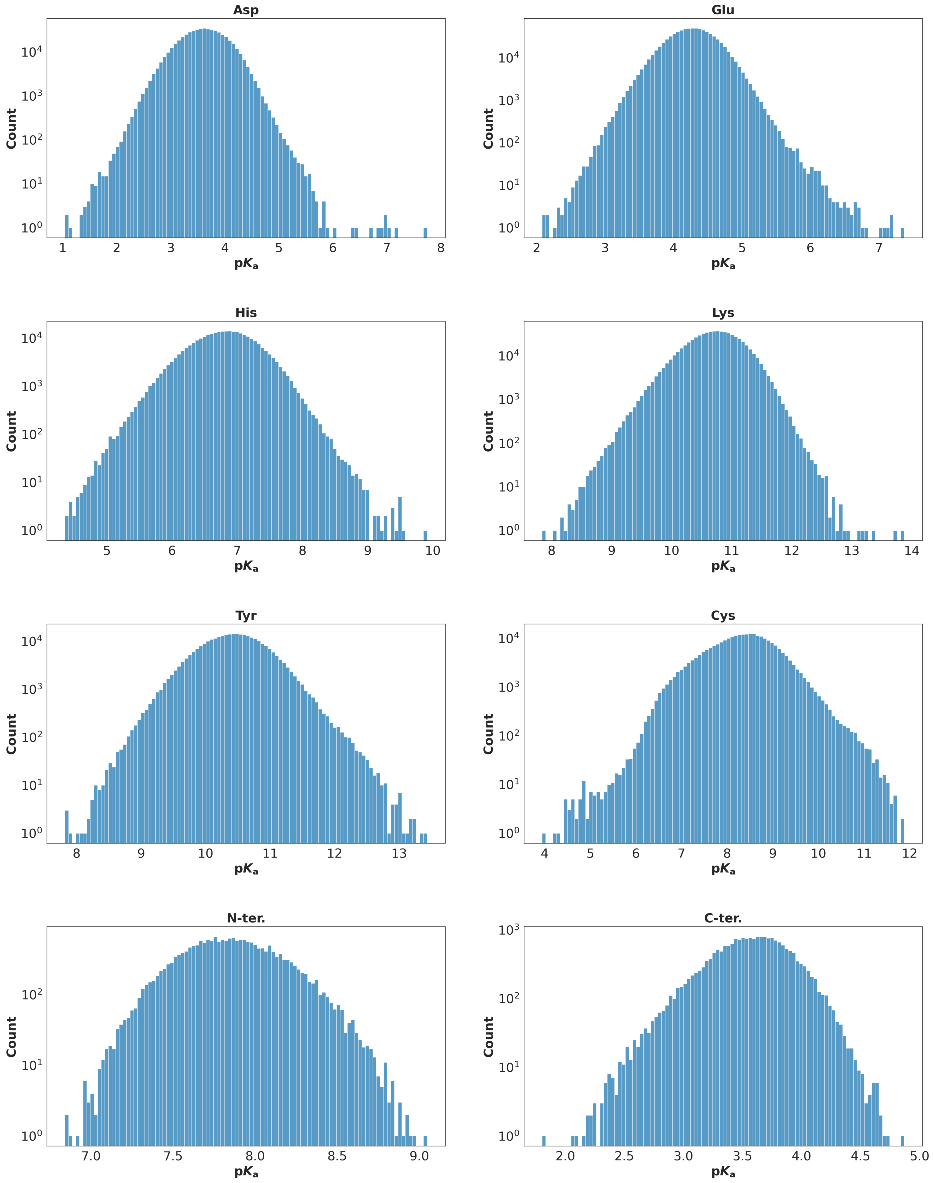


Figure S. 6: Distribution of predicted p$K_\mathrm{a}$ values in the human proteome for different residues

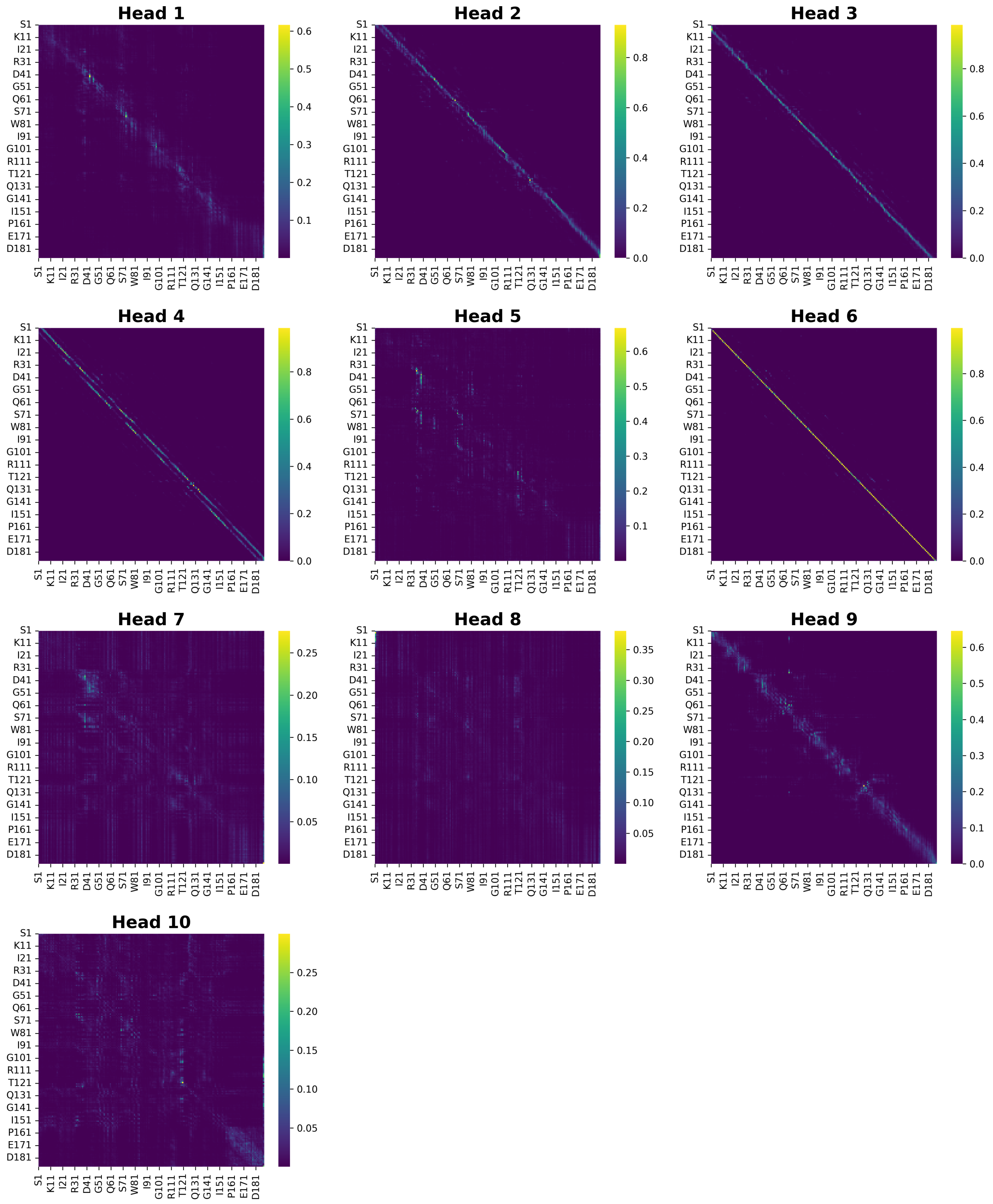


Figure S. 7: The attention maps of from the last layer of ESM2_35M model for AhpC protein (head 1-10)

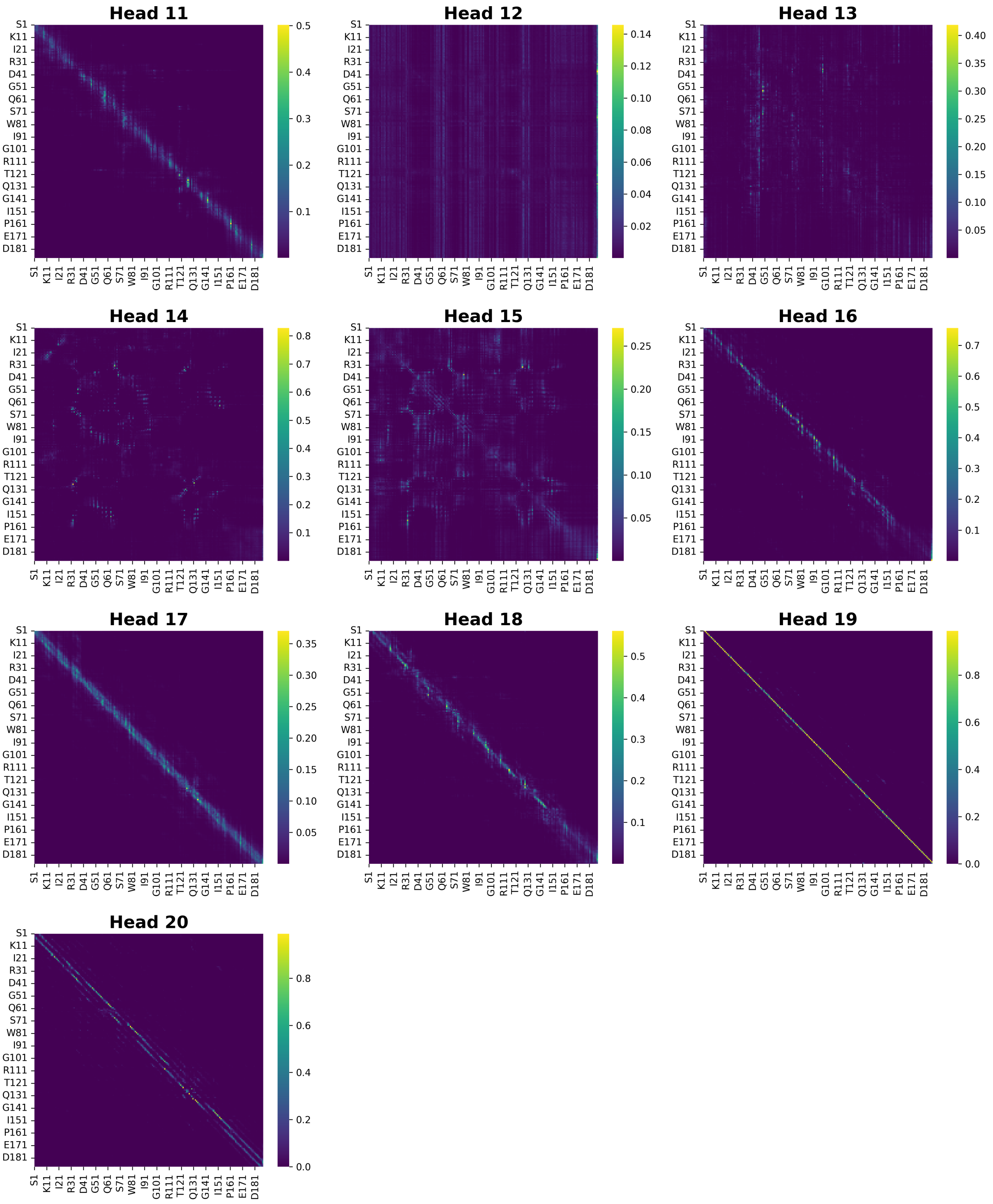


Figure S. 8: The attention maps of the last layer of ESM2_35M model for AhpC protein (head 11-20)

### Supplementary tables

Table S. 1: The information of protein language model used in this study

| Model | Abbr. | # layers | # params | Dim. | Data set |
|---|---|---|---|---|---|
| esm1b_t33_650M_UR50S | ESM1B_650M | 33 | 650M | 1280 | UR50/S |
| esm2_t6_8M_UR50D | ESM2_8M | 6 | 8M | 320 | UR50/D |
| esm2_t12_35M_UR50D | ESM2_35M | 12 | 35M | 480 | UR50/D |
| esm2_t30_150M_UR50D | ESM2_150M | 30 | 150M | 640 | UR50/D |
| esm2_t33_650M_UR50D | ESM2_650M | 33 | 650M | 1280 | UR50/D |
| esm2_t36_3B_UR50D | ESM2_3B | 36 | 3B | 2560 | UR50/D |
| esm2_t48_15B_UR50D | ESM2_15B | 48 | 15B | 5120 | UR50/D |
| prot_t5_xl_bfd | PT_BFD | 24 | ∼3B | 1024 | BFD |
| prot_t5_xl_uniref50 | PT_UR | 24 | ∼3B | 1024 | UR50 |

Table S. 2: The standard p$K_\mathrm{a}$ for different residues

| Residue | Standard p$K_\mathrm{a}$ |
|---|---|
| Arg | 12.5 |
| Glu | 4.5 |
| Lys | 10.5 |
| Cys | 9.0 |
| His | 6.5 |
| Asp | 3.8 |
| Tyr | 10.0 |
| C-ter. | 3.2 |
| N-ter. | 8.0 |

Table S. 3: The tuned hyperparameters for the pKALM model and pI models

| Model | Hidden state dimension | Number of layers |
|---|---|---|
| pKALM | [64, 128, 256, 512] | [1, 2, 3, 4] |
| peptide pI model | [16, 32, 64, 128] | [1, 2, 3, 4, 5] |
| protein pI model | [32, 64, 128, 256] | [1, 2, 3, 4, 5] |

Table S. 4: The comparison of test RMSE of different models for different residues.

| Method | Asp | Glu | His | Lys | Cys | Tyr | C-ter. | N-ter. |
|---|---|---|---|---|---|---|---|---|
| PypKa | 1.2130 | 0.8499 | 1.3601 | 1.1436 | 3.3835 | 1.1068 | 0.7496 | 1.2390 |
| PropKa | 1.3966 | 0.9574 | 1.1254 | 1.0058 | 5.2693 | 0.9896 | / | / |
| PKAI | 1.2659 | 0.9040 | 1.3081 | 1.0515 | 5.1371 | 1.2156 | / | / |
| PKAI+ | 1.1244 | 0.6662 | 1.0809 | 1.0347 | 4.2194 | 1.0679 | / | / |
| DeepKa | 1.1292 | 0.7014 | 1.1848 | 0.9522 | / | / | / | / |
| pKALM | 0.6929 | 0.6641 | 1.0468 | 0.7701 | 2.0984 | 0.6479 | 0.8038 | 0.7419 |
| −pipep | 0.7066 | 0.6615 | 1.0565 | 0.7905 | 2.0854 | 0.6127 | 0.7765 | 0.7208 |
| −piprot | 0.7432 | 0.6580 | 1.0820 | 0.7959 | 1.8850 | 0.6891 | 0.8171 | 0.6382 |
| −pp | 0.7297 | 0.6317 | 1.0726 | 0.8030 | 1.8866 | 0.6697 | 0.8131 | 0.7100 |

* The rows "−pipep", "−piprot", and "−pp" indicate the performance of the pKALM models without peptide pI models, without protein pI models, and without both models, respectively.

Table S. 5: The comparison of test MAE of different models for different residues.

| Method | Asp | Glu | His | Lys | Cys | Tyr | C-ter. | N-ter. |
|---|---|---|---|---|---|---|---|---|
| PypKa | 0.7755 | 0.6515 | 0.9030 | 0.9354 | 3.3827 | 0.9904 | 0.6542 | 0.9180 |
| PropKa | 0.8863 | 0.7265 | 0.8865 | 0.8193 | 4.9575 | 0.6833 | / | / |
| PKAI | 0.7545 | 0.6350 | 0.9365 | 0.8015 | 4.8325 | 1.1533 | / | / |
| PKAI+ | 0.6511 | 0.4902 | 0.7665 | 0.7980 | 3.8975 | 0.8983 | / | / |
| DeepKa | 0.7530 | 0.5306 | 0.7924 | 0.6039 | / | / | / | / |
| pKALM | 0.4730 | 0.4702 | 0.7133 | 0.5625 | 1.7706 | 0.4415 | 0.6783 | 0.5745 |
| −pipep | 0.4957 | 0.4543 | 0.7432 | 0.5812 | 1.7462 | 0.4020 | 0.6404 | 0.6266 |
| −piprot | 0.5219 | 0.4595 | 0.7391 | 0.5578 | 1.5620 | 0.4589 | 0.6508 | 0.5278 |
| −pp | 0.5054 | 0.4486 | 0.7550 | 0.5868 | 1.5867 | 0.4319 | 0.6564 | 0.6150 |

* The rows "−pipep", "−piprot", and "−pp" indicate the performance of the pKALM models without peptide pI models, without protein pI models, and without both models, respectively.

Table S. 6: The comparison of test RMSE of different models for different shift ranges, buried/exposed, and moajor/total residues.

| Method | 0-0.5 | 0.5-1.0 | 1.0-1.5 | 1.5-2.0 | 2.0- | Buried | Exposed | Abundant | Total |
|---|---|---|---|---|---|---|---|---|---|
| PypKa | 0.6048 | 0.7703 | 1.4103 | 1.6671 | 4.2418 | 1.6453 | 0.8032 | 1.1340 | 1.1936 |
| PropKa | 0.6167 | 0.6986 | 1.4145 | 1.4181 | 1.6909 | 2.1226 | 0.9337 | 1.1617 | 1.4507 |
| PKAI | 0.6015 | 0.8084 | 1.7266 | 2.0137 | 4.4972 | 2.1368 | 0.7266 | 1.1345 | 1.4168 |
| PKAI+ | 0.3953 | 0.6834 | 1.4651 | 1.8030 | 4.0603 | 1.7972 | 0.5862 | 0.9631 | 1.1868 |
| DeepKa | 0.4699 | 0.7924 | 1.3643 | 1.1713 | 2.5462 | 1.3534 | 0.7036 | 0.9959 | 0.9959 |
| pKALM | 0.3928 | 0.7156 | 0.9439 | 1.5716 | 1.9515 | 1.1580 | 0.6281 | 0.7740 | 0.8321 |
| −pipep | 0.3748 | 0.6980 | 0.9424 | 1.8229 | 1.8591 | 1.1682 | 0.6215 | 0.7822 | 0.8355 |
| −piprot | 0.3613 | 0.6866 | 0.9352 | 1.6556 | 2.0967 | 1.1722 | 0.6165 | 0.7995 | 0.8392 |
| −pp | 0.4017 | 0.6496 | 0.9549 | 1.6373 | 2.0386 | 1.1434 | 0.6295 | 0.7860 | 0.8288 |

* The rows "−pipep", "−piprot", and "−pp" indicate the performance of the pKALM models without peptide pI models, without protein pI models, and without both models, respectively.

Table S. 7: The comparison of test MAE of different models for different shift ranges, buried/exposed, and moajor/total residues.

| Method | 0-0.5 | 0.5-1.0 | 1.0-1.5 | 1.5-2.0 | 2.0- | Buried | Exposed | Abundant | Total |
|---|---|---|---|---|---|---|---|---|---|
| PypKa | 0.4719 | 0.7045 | 1.2493 | 1.5622 | 4.0462 | 1.2108 | 0.6134 | 0.7792 | 0.8294 |
| PropKa | 0.4365 | 0.6038 | 1.2937 | 1.2925 | 1.6650 | 1.3886 | 0.6928 | 0.8243 | 0.9388 |
| PKAI | 0.4549 | 0.6585 | 1.5575 | 1.7850 | 4.2850 | 1.4359 | 0.5501 | 0.7515 | 0.8844 |
| PKAI+ | 0.3077 | 0.6015 | 1.3913 | 1.7550 | 3.9950 | 1.2387 | 0.4277 | 0.6322 | 0.7347 |
| DeepKa | 0.3607 | 0.6545 | 1.2728 | 1.0341 | 2.4758 | 0.8363 | 0.5223 | 0.6682 | 0.6682 |
| pKALM | 0.2726 | 0.5821 | 0.8367 | 1.5696 | 1.8849 | 0.8126 | 0.4551 | 0.5291 | 0.5644 |
| −pipep | 0.2474 | 0.5660 | 0.8045 | 1.8224 | 1.7903 | 0.8278 | 0.4553 | 0.5388 | 0.5708 |
| −piprot | 0.2465 | 0.5587 | 0.7818 | 1.6538 | 2.0343 | 0.8412 | 0.4472 | 0.5459 | 0.5722 |
| −pp | 0.2678 | 0.5370 | 0.8132 | 1.6209 | 1.9923 | 0.8074 | 0.4671 | 0.5429 | 0.5716 |

* The rows "−pipep", "−piprot", and "−pp" indicate the performance of the pKALM models without peptide pI models, without protein pI models, and without both models, respectively.
